# The m^6^A methyltransferase complex protein VIRMA engages hepatitis C virus RNA through a WTAP-independent mechanism

**DOI:** 10.64898/2026.09.11.750944

**Authors:** Katherine M. Bland, Jordan V. Reaves, Yechan Moon, Yue Deng, Moonhee Park, Stacy M. Horner

**Affiliations:** Department of Molecular Genetics and Microbiology, Duke University School of Medicine, Durham, NC 27710, USA; Program in Cell and Molecular Biology, Duke University School of Medicine, Durham, NC 27710, USA; Department of Integrative Immunobiology, Duke University School of Medicine, Durham, NC 27710, USA; Department of Medicine, Duke University School of Medicine, Durham, NC 27710, USA

**Keywords:** N6-methyladenosine, m6A, VIRMA, KIAA1429, WTAP, HCV, RNA modification

## Abstract

Cellular proteins in the m^6^A methyltransferase protein complex (m^6^A-MTC) work together to modify both cellular and viral RNAs with N6-methyladenosine (m^6^A), an RNA modification that regulates viral infection. We previously showed that hepatitis C virus (HCV), a cytoplasmic positive-sense RNA virus, has an m^6^A-modified RNA genome. We also found that m^6^A deposition on HCV RNA requires WTAP, which promotes association of the METTL3-METTL14 catalytic heterodimer to the viral RNA. However, the factors that directly recognize HCV RNA and recruit the m^6^A-MTC to the viral genome are unknown. Here, we identify the cellular m^6^A-MTC component VIRMA as an HCV RNA-targeting factor required for METTL3 association with the viral genome. We show that endogenous VIRMA can associate with both HCV and cellular RNAs independently of WTAP. Further, we define distinct domains of VIRMA involved in WTAP interaction and RNA binding and show that the regions of VIRMA involved in WTAP-interaction are required for cellular but not HCV RNA association. Together, these results establish that VIRMA uses distinct molecular determinants to engage viral and cellular RNAs and define VIRMA as a substrate-recognition factor required for m^6^A-MTC association with HCV RNA.

## INTRODUCTION

Hepatitis C virus (HCV) is a positive-sense, single-stranded RNA virus of the *Flaviviridae* family that causes substantial global morbidity and mortality. As of 2024, an estimated 47 million people were living with HCV infection, with approximately 0.9 million new infections and 240,000 HCV-related deaths reported worldwide (World Health Organization 2026). HCV uses its genome directly as messenger RNA to produce a single polyprotein, encodes its own RNA-dependent RNA polymerase NS5B to replicate its genome, and carries out translation, RNA replication, and particle assembly in the cytoplasm on ER-associated membranes (Lindenbach and Rice 2005; Romero-Brey et al. 2012). Beyond these core features, the HCV genome is extensively regulated at the RNA level: it is stabilized by the liver-specific microRNA miR-122 (Sarnow and Sagan 2016), translated via a structured internal ribosomal entry site in its 5′ UTR (Filbin and Kieft 2009), and dynamically bound by cellular RNA-binding proteins that regulate viral RNA translation, replication, and stability (Sarnow and Sagan 2016). More recently, it has become clear that the genomes of positive-sense, single-stranded RNA viruses, including HCV, can contain RNA modifications, such as N6-methyladenosine (m^6^A), which adds a further layer of RNA regulation to viral infection (Gokhale et al. 2016; Baquero-Perez et al. 2021; Horner and Reaves 2024).

The RNA modification m^6^A can be dynamically added to both viral and cellular RNAs during infection. In host cells, m^6^A is deposited co-transcriptionally in the nucleus (Slobodin et al. 2017) by a multiprotein methyltransferase complex (m^6^A-MTC). The m^6^A-MTC contains the METTL3-METTL14 catalytic heterodimer and associated regulatory factors, including WTAP, VIRMA, ZC3H13, and RBM15-RBM15B that influence its assembly, transcript-selective methylation, and localization (Liu et al. 2014; Schwartz et al. 2014; Garcias Morales and Reyes 2021; Su et al. 2022). The functional consequences of m^6^A addition or removal arise largely from changes in RNA binding protein interactions (Gilbert and Nachtergaele 2023). These RNA binding proteins, commonly called m^6^A-readers, can dictate whether an mRNA molecule is degraded or translated, ultimately influencing gene expression (Wang et al. 2014; Wang et al. 2015; Arguello et al. 2017; Edupuganti et al. 2017). On viral RNA, we have shown that HCV RNA is m^6^A modified and that m^6^A within the E1 coding region negatively regulates infectious virion production in a METTL3 and METTL14-dependent manner (Gokhale et al. 2016). Additionally, depletion of m^6^A machinery components has been implicated in HCV IRES-mediated translation and in modulating RIG-I sensing of HCV RNA (Kim et al. 2020; Kim and Siddiqui 2021). WTAP, an m^6^A-MTC component that scaffolds METTL3-METTL14 and directs their recruitment to RNA substrates (Ping et al. 2014), is also required for m^6^A modification of HCV RNA (Sacco et al. 2022). Interestingly, we also found that a WTAP mutant lacking its nuclear localization signal is sufficient to direct METTL3 and METTL14 to methylate HCV RNA but not cellular RNAs, revealing a non-canonical mechanism for m^6^A-MTC targeting to viral RNA (Sacco et al. 2022). However, the factor(s) that directly recognize HCV RNA and recruit the m^6^A-MTC to the viral genome remain unknown.

VIRMA, also referred to as KIAA1429, is a strong candidate for the RNA-targeting factor that recruits the m^6^A-MTC to HCV RNA. VIRMA is the largest subunit of the m^6^A-MTC (Yue et al. 2018; Su et al. 2022) and directly interacts with WTAP to form the structural core of the complex (Bawankar et al. 2021; Su et al. 2022; Yan et al. 2022). VIRMA is an RNA-binding protein that associates with cellular transcripts and directs m^6^A deposition to specific transcripts and transcript regions, including 3′ UTRs and near stop codons (Yue et al. 2018). Although VIRMA is predominantly nuclear at steady state (Yue et al. 2018; Lee et al. 2023), cytosolic functions of VIRMA have been reported (Li et al. 2023). In particular, VIRMA was previously identified as an interactor of several non-nuclear HCV proteins in a systematic proteomic screen of HCV-host protein interactions (Ramage et al. 2015). Although a small number of studies have implicated VIRMA in viral infection (Srinivas et al. 2021; Zhang et al. 2021; N’Da Konan et al. 2022), whether VIRMA has an analogous RNA-targeting function on viral RNAs has not been directly tested.

Here, we identify VIRMA as the substrate-recognition factor for the m^6^A-MTC on HCV RNA. We show that VIRMA promotes METTL3 association with HCV RNA and contributes to m^6^A deposition on the viral genome. Endogenous VIRMA can bind both cellular and HCV RNAs independently of its interaction with WTAP. However, the regions of VIRMA involved in WTAP-interaction are required for binding to cellular but not HCV RNA. These findings identify HCV and cellular RNAs as distinct classes of VIRMA substrates and establish VIRMA as a substrate-recognition factor that directs the m^6^A-MTC to HCV RNA.

## RESULTS

### VIRMA is essential for m^6^A modification of HCV RNA

To investigate the role of VIRMA in m^6^A modification of HCV RNA, we performed m^6^A-methylated RNA-immunoprecipitation followed by qPCR (meRIP-qPCR) of RNA harvested from HCV-infected Huh7 cells (MOI 0.3, 48 h) that had been treated with a VIRMA-targeting siRNA or a non-targeting control siRNA. The specificity of the meRIP assay was validated by inclusion of unmodified and m^6^A-modified spike-in RNAs (**Figure S1**). VIRMA depletion reduced m^6^A levels on HCV RNA by ∼75% and on the cellular m^6^A-modified transcript *ACTB* by ∼60% (**Figure 1A-B**). Depletion of METTL3-METTL14 or WTAP also reduced m^6^A on both HCV RNA and on *ACTB,* consistent with our previous findings (Gokhale et al. 2016; Sacco et al. 2022).

**Figure 1:**
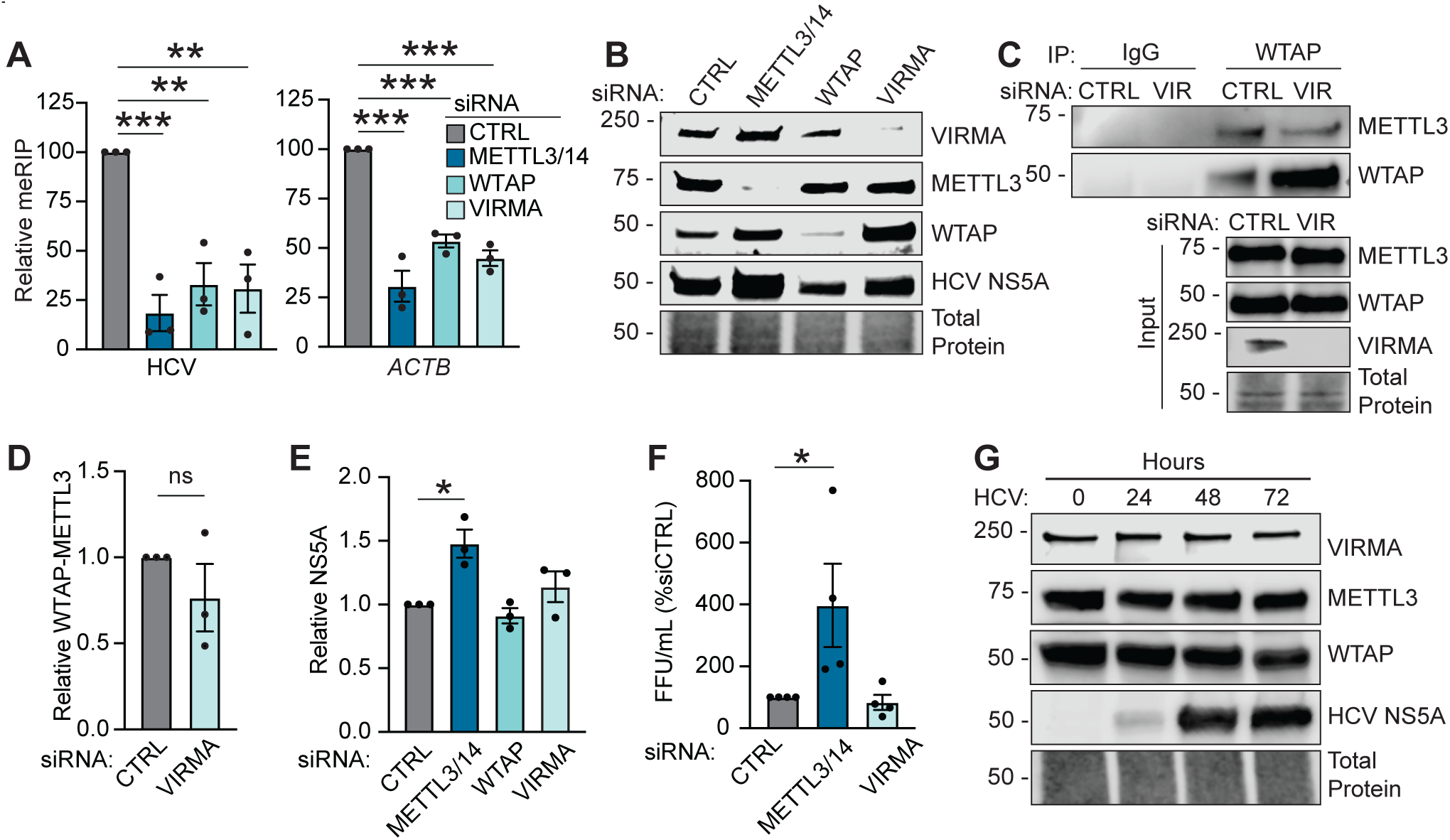
VIRMA is essential for m⁶A modification of HCV RNA. **(A)** Relative m^6^A enrichment of HCV and *ACTB* in Huh7 cells treated with the indicated siRNAs and infected with HCV (MOI 0.3, 48 h). The results are displayed as the percentage of the input for each condition, normalized to siCTRL. **(B)** Immunoblot analysis of HCV-infected (MOI 0.3, 48 h) Huh7 cells treated with indicated siRNAs. **(C)** Endogenous WTAP immunoprecipitation and immunoblot analysis of WTAP and METTL3 from Huh7 cells treated with siCTRL or siVIRMA, with IgG as an isotype control. (**D**) Quantification showing relative WTAP-METTL3 interaction (from C), normalized to total protein, WTAP input, and siCTRL. **(E)** Quantification showing relative HCV NS5A levels (from B), normalized to total protein and siCTRL. **(F)** Infectious HCV titers measured by focus-forming assay from supernatants of Huh7 cells treated with the indicated siRNA and infected with HCV (MOI 0.3, 48 h). **(G)** Immunoblot analysis during a 72-hour time course of HCV infection (MOI 0.3). Graphs show the mean +/- SEM (n=3 or 4 (for F) biological replicates, as indicated by individual replicate data points), analyzed by one-way ANOVA with Sidak’s multiple-comparison test or Welch’s test (for D) (* - P<0.05, ** - P<0.01, *** - P<0.001).

We next asked whether the loss of m^6^A upon VIRMA depletion could be explained indirectly, through loss or disruption of the m^6^A-MTC. Because prior studies reported that VIRMA depletion destabilizes WTAP protein (Yue et al. 2018; Bawankar et al. 2021), and reduced WTAP levels alone would decrease m^6^A on HCV RNA (Sacco et al. 2022), we measured WTAP protein levels by immunoblot. In HCV-infected Huh7 cells, WTAP protein levels were not reduced by siVIRMA (**Figure 1B**), ruling out reduced WTAP as an explanation for the m^6^A loss. As VIRMA can also stabilize the m^6^A-MTC (Su et al. 2022), we next tested whether VIRMA depletion disrupts the WTAP-METTL3 interaction by performing an endogenous WTAP-METTL3 co-immunoprecipitation in the presence or absence of VIRMA. Endogenous WTAP-METTL3 co-immunoprecipitation, along with quantification, revealed that this interaction is preserved in the absence of VIRMA (**Figure 1C-1D**). Together, these data indicate that VIRMA is required for m^6^A modification of HCV RNA independently of changes in WTAP protein levels or WTAP-METTL3 interaction.

Finally, we asked how VIRMA depletion compared to depletion of METTL3-METTL14 or WTAP at the level of HCV infection. VIRMA depletion did not alter HCV NS5A protein levels (**Figure 1E**) or infectious virus in the supernatant (**Figure 1F**). Consistent with our previous work, METTL3-METTL14 depletion increased both NS5A levels and infectious virus in the supernatant (**Figure E-1F**) (Gokhale et al. 2016), while WTAP depletion decreased NS5A levels (**Figure 1E**) (Sacco et al. 2022). Immunoblot analysis across a time course of HCV infection showed no apparent change in VIRMA, WTAP, or METTL3 protein levels **(Figure 1G**), indicating that HCV infection does not alter the abundance of the m^6^A-MTC components involved in HCV RNA methylation. Together, these data establish that VIRMA is required for m^6^A modification of both HCV RNA and the host transcript *ACTB*, but that VIRMA depletion produced a distinct effect on HCV infection despite loss of m^6^A.

### VIRMA is required for METTL3 association with HCV RNA

Within the m^6^A-MTC, VIRMA is connected to METTL3 primarily through interactions with WTAP (Su et al. 2022; Yan et al. 2022). Because WTAP mediates METTL3 binding to HCV RNA (Sacco et al. 2022), we asked whether VIRMA is similarly required for METTL3 binding to HCV RNA. To do this, we immunoprecipitated endogenous METTL3 from HCV-infected Huh7 cells (MOI 1, 48 h) following depletion of VIRMA, WTAP (as a positive control), or a non-targeting control siRNA, and confirmed comparable METTL3 pulldown across conditions by immunoblot (**Figure 2A**). RT-qPCR analysis of METTL3-bound RNA revealed that VIRMA depletion reduced METTL3 binding to HCV RNA to an extent comparable to WTAP depletion, and similarly reduced METTL3 binding to the methylated cellular transcripts *ACTB* and *SON* (**Figure 2B**). Together, these data demonstrate that VIRMA is required for METTL3 binding to both HCV RNA and cellular transcripts, consistent with a VIRMA-WTAP-METTL3 recruitment axis established by prior structural and biochemical studies (Yue et al. 2018; Bawankar et al. 2021; Su et al. 2022; Yan et al. 2022).

**Figure 2:**
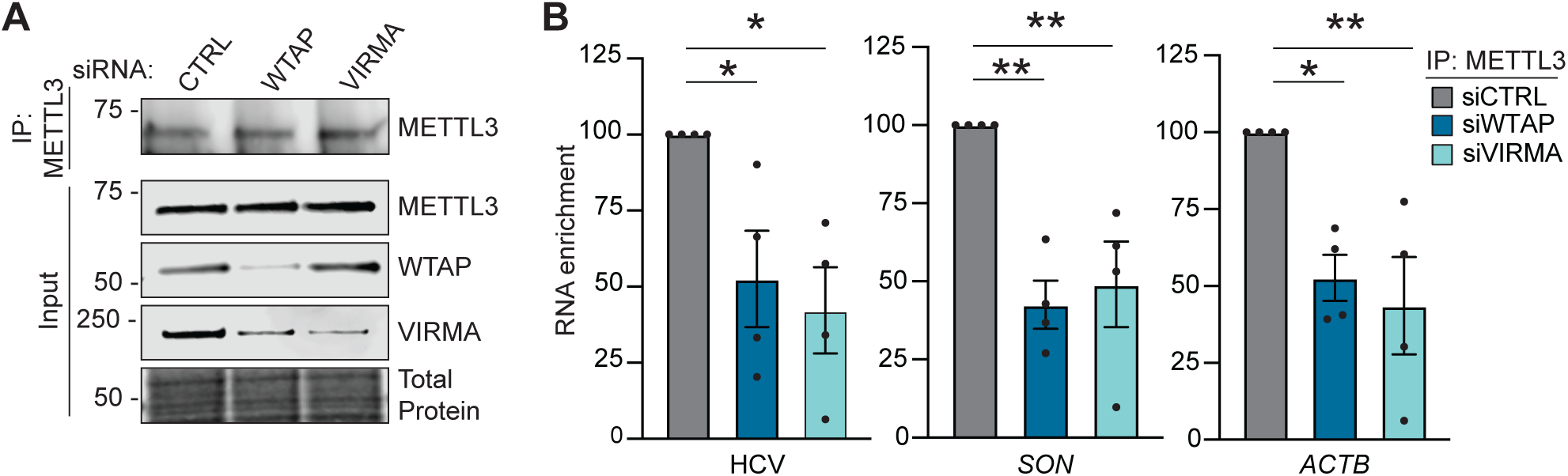
VIRMA is required for METTL3 association with HCV RNA. **(A)** Immunoblot analysis of endogenous METTL3 immunoprecipitation from Huh7 cells treated with the indicated siRNA and infected with HCV (MOI 1, 48 h). **(B)** RNA enrichment of HCV RNA and the cellular transcripts *ACTB* and *SON* following METTL3 immunoprecipitation from the samples in panel A. The results are displayed as the percentage of the input for each condition, normalized to siCTRL (set at 100%). Graphs show the mean +/- SEM, n=4 biological replicates, analyzed by one-way ANOVA with Sidak’s multiple-comparison test (* - P<0.05, ** - P<0.01).

### VIRMA binds HCV and cellular RNAs independently of WTAP expression

Having established that VIRMA and WTAP both promote METTL3 targeting to HCV RNA, we next tested whether VIRMA engages HCV RNA in the absence of WTAP. We performed a native RNA immunoprecipitation of endogenous VIRMA in the presence or absence of WTAP. VIRMA-RNA complexes were captured from HCV-infected Huh7 cells (MOI 1, 48 h) treated with siWTAP or a non-targeting control siRNA, using an anti-VIRMA antibody or an IgG control (**Figure 3A)**. VIRMA immunoprecipitation enriched HCV RNA and the cellular transcripts *ACTB* and *SON* over IgG control. Importantly, this binding was preserved following WTAP depletion (**Figure 3B**). Thus, VIRMA-RNA association does not require WTAP, indicating that VIRMA can engage HCV RNA and some cellular RNAs independently of WTAP expression.

**Figure 3:**
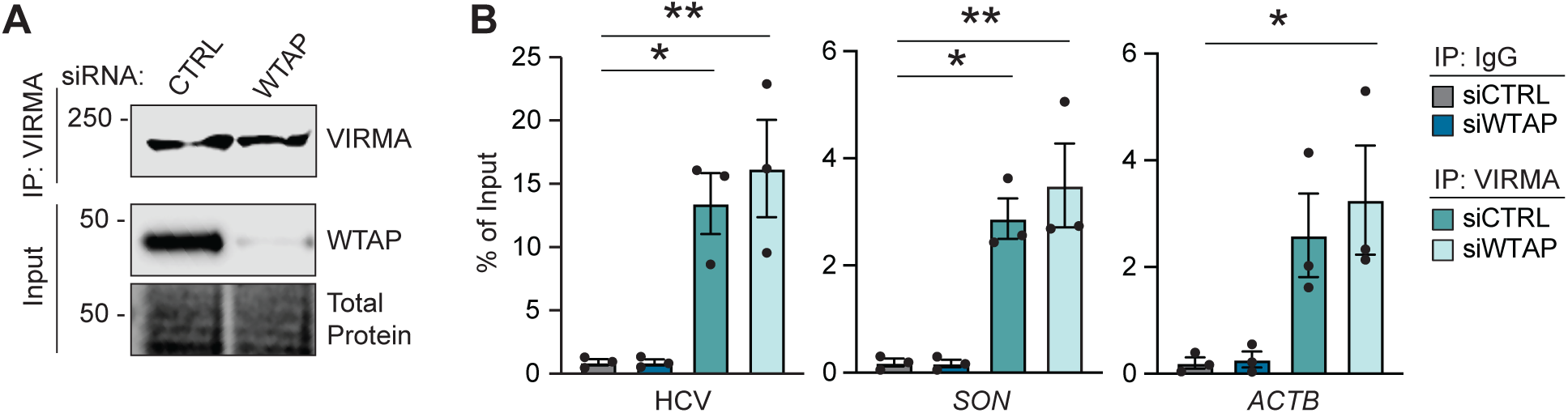
VIRMA binds HCV and cellular RNAs independently of WTAP expression. **(A)** Immunoblot analysis of endogenous VIRMA immunoprecipitation from Huh7 cells treated with siCTRL or siWTAP and infected with HCV (MOI 1, 48 h). **(B)** RNA enrichment of HCV RNA and the cellular transcripts *ACTB* and *SON* following VIRMA or IgG immunoprecipitation, matching the conditions in panel A. The results are displayed as the percentage of the input for each condition. Graphs show the mean +/- SEM, n=3 biological replicates. Data were analyzed by one-way ANOVA with Sidak’s multiple-comparison test (* - P<0.05, ** - P<0.01).

### VIRMA regions contribute differentially to HCV and cellular RNA engagement

To separate the RNA binding activity of VIRMA from its scaffolding role in the m^6^A-MTC, we generated FLAG-VIRMA constructs lacking regions implicated in WTAP interaction or RNA binding **(Figure 4A)**. Cryo-EM structures of the human m^6^A-MTC show that VIRMA contacts the WTAP homodimer through two spatially distinct interfaces, aa 335-526 (WTAP-a) and aa 943-1147 (WTAP-b) (Su et al. 2022; Yan et al. 2022). To disrupt these interfaces, we generated VIRMA^ΔWTAP^ in which both WTAP-a and WTAP-b were deleted (**Figure 4A**). Because WTAP mediates the connection of VIRMA to METTL3 within the m^6^A-MTC, VIRMA^ΔWTAP^ is predicted to disrupt the association of VIRMA with the WTAP-METTL3 complex while leaving the rest of the protein intact. To disrupt RNA binding, we drew on the published protein-RNA crosslinking data that identified VIRMA-RNA crosslinks at P140, E1597, and F1706, and on functional data showing that truncation of VIRMA at aa 1586 reduces RNA binding by the reconstituted m^6^A-MTC *in vitro* (Su et al. 2022). Based on these findings, we generated VIRMA^ΔRBD^, which lacks aa 1586-1812 **(Figure 4A)**.

**Figure 4:**
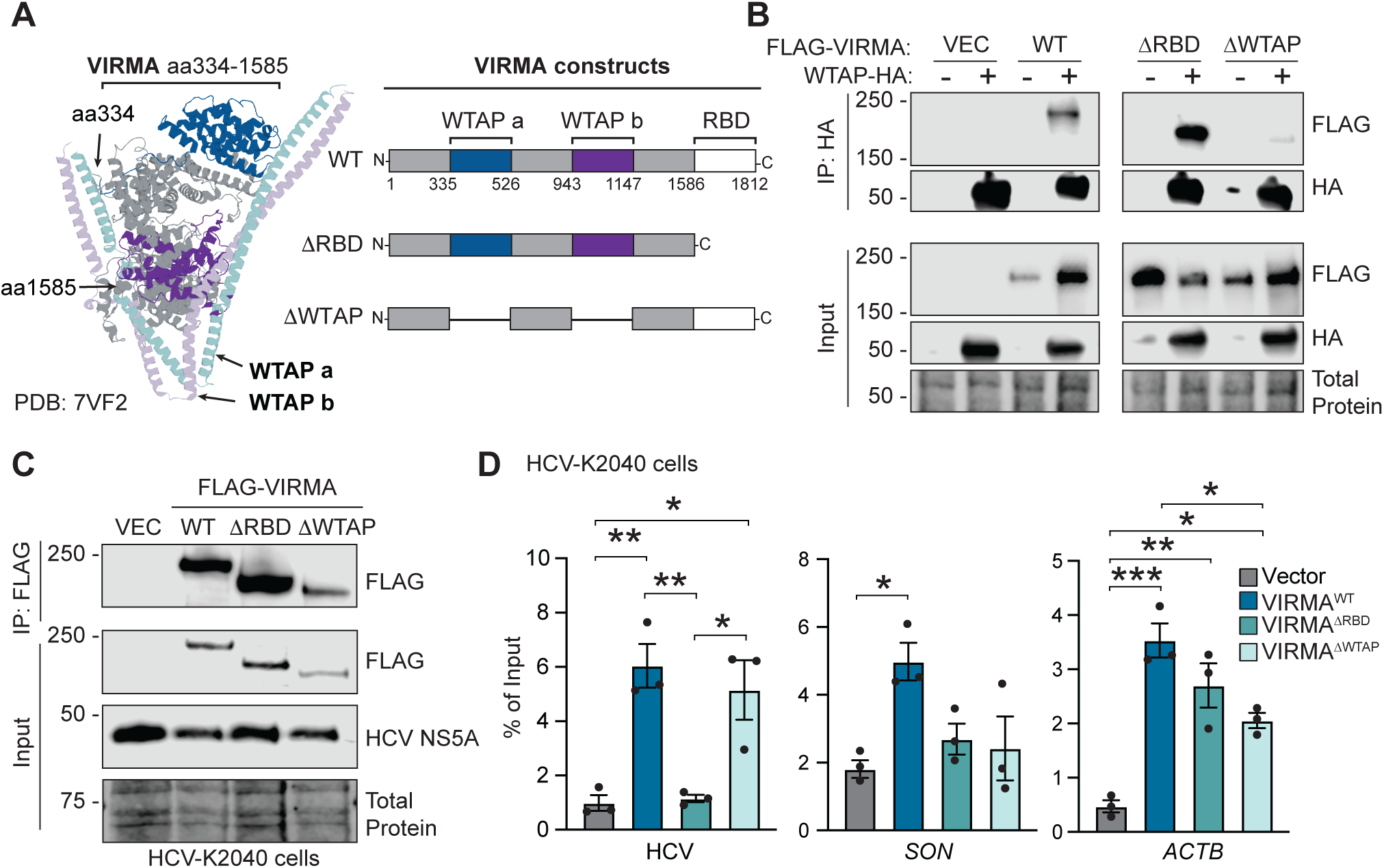
VIRMA regions contribute differentially to HCV and cellular RNA engagement. **(A)** Structural model of the human m^6^A-MTC (PDB: 7VF2), showing VIRMA aa 334-1585 and the WTAP homodimer, as well as the regions of VIRMA involved in these contacts. The C-terminal RNA binding region was not captured in this model. To the right, schematic representations of the FLAG-tagged VIRMA wild-type (WT), ΔRBD, and ΔWTAP constructs are shown. **(B)** Co-immunoprecipitation of WTAP-HA with FLAG-tagged VIRMA^WT^, VIRMA^ΔRBD^, or VIRMA^ΔWTAP^ expressed in Huh7 cells. **(C)** Representative immunoblot showing FLAG immunoprecipitation and expression of FLAG-tagged VIRMA constructs in Huh7-K2040 cells that stably replicate HCV subgenomic RNA, as shown by HCV NS5A expression in the immunoblot. **(D)** RNA enrichment of HCV RNA and the cellular transcripts *SON* and *ACTB* from samples in panel C following FLAG immunoprecipitation. The results are displayed as the percentage of the input for each condition. Immunoblots are representative of n=3 biological replicates. Graphs show the mean +/- SEM, n=3 biological replicates. Data were analyzed by one-way ANOVA with Sidak’s multiple-comparison test (* - P<0.05, ** - P<0.01, *** - P<0.001).

Co-immunoprecipitation of WTAP-HA with FLAG-VIRMA revealed that VIRMA^ΔWTAP^ failed to bind WTAP, while VIRMA^ΔRBD^ retained WTAP interaction comparable to VIRMA^WT^ (**Figure 4B**). The preserved WTAP interaction of VIRMA^ΔRBD^ indicates that the C-terminal region of VIRMA is dispensable for WTAP binding in cells, despite an earlier biochemical study identifying a C-terminal VIRMA fragment (aa 1575-1812) as WTAP-interacting (Bawankar et al. 2021). Together, these results isolate the WTAP-interacting and RNA-binding regions as functionally separable in cells.

To determine how these VIRMA regions contribute to viral and cellular RNA engagement, we used cells harboring a stably replicating HCV subgenomic RNA (Huh7-K2040) (Sumpter et al. 2004), as stable expression of VIRMA could not be achieved in Huh7 cells. We transiently expressed empty vector, FLAG-VIRMA^WT^, FLAG-VIRMA^ΔRBD^ and FLAG-VIRMA^ΔWTAP^ in Huh7-K2040 cells. All VIRMA constructs displayed similar subcellular localization (**Figure S2**), indicating that deletion of these regions did not grossly alter localization. We then performed a FLAG-RNA immunoprecipitation **(Figure 4C)** followed by RT-qPCR for HCV RNA and the m^6^A-modified cellular transcripts *SON* and *ACTB* (**Figure 4D**). Deletion of the C-terminal RNA-binding region reduced enrichment of HCV RNA and *SON* to levels comparable to vector control, whereas *ACTB* retained significant enrichment above vector (**Figure 4D**). Thus, while the C-terminal region is required for VIRMA to bind HCV RNA and *SON,* it only partially contributes to *ACTB* association. Deletion of the WTAP-interaction regions had a distinct pattern: VIRMA^ΔWTAP^ showed reduced *SON* enrichment and reduced but still significant *ACTB* enrichment **(Figure 4D)**, whereas VIRMA^ΔWTAP^ associated with HCV RNA at levels comparable to VIRMA^WT^, indicating that the WTAP-interaction regions are dispensable for HCV RNA engagement. Notably, this requirement was independent of WTAP protein, as WTAP depletion did not reduce VIRMA-cellular RNA binding **(Figure 3)**. Together, these data reveal distinct requirements for VIRMA engagement of viral and some cellular RNAs. HCV RNA binding requires the C-terminal RNA-binding region but not the WTAP-interaction regions or WTAP protein. In contrast, cellular RNA engagement shows transcript-specific dependence on both the C-terminal region and the WTAP-interaction regions.

## DISCUSSION

Previous work has established that METTL3, METTL14, and WTAP are required for m^6^A modification of HCV RNA, but the RNA-targeting factor that promotes methylation of viral RNA has remained unknown. Here, we identify VIRMA as a substrate recognition factor that promotes recruitment of the m^6^A-MTC to HCV RNA. Depletion of VIRMA reduced m^6^A modification and METTL3 association with the viral RNA. VIRMA association with HCV RNA and the cellular RNAs tested was maintained following WTAP depletion, demonstrating that VIRMA can engage these RNA substrates independently of WTAP. Deletion of the RNA-binding domain of VIRMA reduced engagement of both HCV and cellular RNAs, whereas deletion of the WTAP-interaction region impaired association with cellular RNAs but not HCV RNA.

The ability of VIRMA to associate with RNA independently of WTAP has important implications for how the m^6^A-MTC is targeted to its substrates. Within the m^6^A-MTC, WTAP mediates the principal connection between VIRMA and the METTL3-METTL14 catalytic core (Su et al. 2022), implying a model in which VIRMA-WTAP interaction bridges RNA substrate recognition to catalysis. However, whether this interaction is required for VIRMA to engage RNA has remained unclear. Endogenous VIRMA remained associated with both HCV and the cellular RNAs after WTAP loss, indicating that RNA engagement does not require the VIRMA-WTAP interaction. In parallel, VIRMA depletion reduced METTL3 association with these RNAs without disrupting the interaction between WTAP and METTL3. Together, these findings support a model in which VIRMA-mediated RNA engagement is mechanistically separable from its association with the larger m^6^A-MTC and promotes efficient recruitment of METTL3 to RNA substrates. In this architecture, distinct regions of VIRMA mediate RNA substrate binding and MTC association, consistent with a role for VIRMA as an RNA adaptor for the m^6^A-MTC.

Structure-function analysis of VIRMA further separated the requirements for its engagement with HCV RNA and cellular RNAs. Deletion of the previously established RNA-binding domain of VIRMA (Su et al. 2022) reduced enrichment of both HCV RNA and cellular RNAs without disrupting WTAP interaction, indicating that this region is required for engagement of both substrate classes. In contrast, deletion of its WTAP-interaction domain impaired association to cellular RNAs but did not affect HCV RNA association. In cells, the C-terminus of VIRMA was dispensable for WTAP interaction, in contrast to an earlier biochemical study that identified this region as WTAP-interacting (Bawankar et al. 2021). This C-terminal WTAP-binding activity may therefore be dispensable in the context of full-length VIRMA, or it may represent an indirect association not captured in cellular co-immunoprecipitation.

Cellular RNA engagement required the WTAP-interaction region of VIRMA but not WTAP protein itself. Endogenous VIRMA retained binding to cellular RNAs after WTAP depletion (**Figure 3**), yet deletion of the WTAP-interaction region reduced cellular RNA binding (**Figure 4D**). One interpretation of this dissociation is that the WTAP-interaction region has a second, WTAP-independent function that contributes to cellular RNA engagement. This function could involve a conformational role important for cellular substrate engagement, direct RNA binding, or interactions with additional RNA-binding cofactors.

The molecular determinants that direct VIRMA specifically to HCV RNA also remain unknown. VIRMA could recognize features of the HCV RNA sequence or structure, including its highly structured IRES (Filbin et al. 2013; Wan et al. 2022), or be recruited through viral or cellular RNA-binding proteins. Indeed, systematic proteomic screens of HCV-host protein interactions have identified VIRMA (KIAA1429) as a potential HCV-interacting protein (Ramage et al. 2015). Whether VIRMA recognizes HCV RNA directly or through additional factors will be an important question for defining how the m^6^A-MTC is targeted to viral RNA.

The mechanism by which VIRMA promotes m^6^A-MTC targeting to HCV RNA is particularly relevant given the exclusively cytoplasmic life cycle of HCV. HCV is a positive-sense, single-stranded RNA virus whose genome serves directly as messenger RNA and as the template for replication, without a DNA intermediate or nuclear phase (Lindenbach and Rice 2005). Current models of cellular m^6^A deposition propose that the m^6^A-MTC is recruited to RNA co-transcriptionally within the nucleus through interactions with RNA polymerase II and associated RNA-binding proteins (Slobodin et al. 2017). HCV, in contrast, encodes its own RNA-dependent RNA polymerase and completes its replication cycle in re-arranged ER-derived membrane compartments in the cytoplasm (Moradpour et al. 2007; Paul and Bartenschlager 2015), placing viral RNA outside this nuclear targeting mechanism. Our previous work provided evidence that HCV m^6^A deposition can occur outside the canonical nuclear pathway: a WTAP mutant lacking its nuclear localization signal fails to support cellular mRNA methylation but retains the ability to support HCV RNA methylation (Sacco et al. 2022). Our finding that VIRMA associates with HCV RNA independently of WTAP suggests that VIRMA-mediated substrate engagement can occur without prior assembly of the canonical nuclear m^6^A-MTC.

Despite reducing m^6^A modification of HCV RNA, VIRMA depletion did not phenocopy loss of other m^6^A-MTC components with respect to HCV protein levels or infectious virion production. Depletion of METTL3-METTL14 increased HCV protein levels and infectious virus production, whereas VIRMA depletion had no significant effect on either phenotype. This phenotypic difference may reflect distinct roles for m^6^A-MTC components in shaping infectious outcomes. Our previous work showed that the effects of m^6^A on HCV infection are mediated by modification of both viral and cellular RNAs, raising the possibility that VIRMA, WTAP, and METTL3-METTL14 differentially regulate host transcripts that influence infection (Gokhale et al. 2020; Sacco et al. 2022). Alternatively, m^6^A modifications within distinct regions of HCV RNA have been associated with both proviral and antiviral effects on infection (Gokhale et al. 2016; Kim et al. 2020; Kim and Siddiqui 2021), such that total HCV RNA methylation may not directly predict how loss of individual m^6^A-MTC components affects infection.

Together, our findings identify VIRMA as a substrate-recognition factor that promotes recruitment of the m^6^A-MTC to HCV RNA and reveal that RNA substrate engagement can be mechanistically separated from canonical m^6^A-MTC assembly. The distinct requirements for VIRMA association with HCV and cellular RNAs suggest that individual RNA transcripts, HCV and cellular, may be recognized by the m^6^A-MTC through different molecular configurations rather than a single unified mechanism. This study provides an example for understanding how the m^6^A-MTC can be directed to a non-nuclear viral RNA and highlights substrate recognition as an important determinant of m^6^A targeting during infection. More broadly, our results identify substrate-recognition factors as a distinct layer of m^6^A regulation and suggest how host m^6^A machinery could be directed to other RNA viruses during infection. Our findings add to a growing body of work showing that RNA viruses reshape host RNA processing by co-opting cellular RNA-binding proteins for functions distinct from their canonical roles (Iselin et al. 2022; Bermudez et al. 2024; Castello and Kamel 2025).

## ACKNOWLEDGEMENTS

We thank those colleagues who generously provided reagents, as indicated in the Materials and Methods; the Duke Functional Genomics Core, and members of the Horner Lab for valuable feedback and discussion. This work was supported by National Institutes of Health grants R01AI125416 (S.M.H) and T32CA00911 and T32GM142605 (K.M.B.).

## AUTHOR CONTRIBUTIONS

Conceptualization, K.M.B and S.M.H.; Methodology, K.M.B and S.M.H.; Formal Analysis, K.M.B and S.M.H.; Investigation, K.M.B, Y.M., J.V.R., Y.D., M.P., and S.M.H.; Writing – Original Draft, K.M.B., and S.M.H.; Writing – Review & Editing, all authors; Supervision, S.M.H.; Funding Acquisition, S.M.H.

## DATA AVAILABILITY

All data generated or analyzed during this study are available within the article and its supplemental information files. Source data for all figures, including individual replicate values for qPCR-based assays and uncropped immunoblot images, will be deposited in a data repository prior to publication. Plasmids generated in this study are available from the corresponding author upon reasonable request.

## METHODS

### Cells and Cell Culture

Cells were grown in Dulbecco’s modification of Eagle’s medium (DMEM; Mediatech) supplemented with 10% fetal bovine serum (FBS) (HyClone), 1X minimum essential medium non-essential amino acids (Thermofisher), and 25 mM HEPES (Thermofisher), referred to as complete DMEM (cDMEM). The identity of the Huh7 and Huh7.5 cells was verified by using the GenePrint STR kit (Duke DNA Analysis Facility). Cells were obtained from the following sources: 293T (CRL-3216; ATCC), Huh7, Huh7.5, and Huh7-K2040 cells (gift of Dr. Michael Gale Jr., University of Washington) (Sumpter et al. 2004; Sumpter et al. 2005). All cell lines were verified as mycoplasma free by the MycoStrip Mycoplasma Detection Kit (InvivoGen).

### Plasmids

The following plasmids were generated by subcloning PCR-generated amplicons from the indicated oligonucleotides into the KpnI-to-PmeI digested pEF-TAK using InFusion recombinase (Takara Bio): pEF-TAK-FLAG-VIRMA^WT^, pEF-TAK-FLAG-VIRMA^ΔRBD^, pEF-TAK-FLAG-VIRMA^ΔWTAP^. Primer sequences for InFusion cloning are listed in Data Table S1. The following additional plasmid was used in this study: pEF-TAK-WTAP-HA (Sacco et al. 2022).

### Antibodies

Antibodies used in this study include: R-anti-VIRMA (1:1000, Proteintech 25712-1-AP), M-anti-VIRMA (1:5000, Proteintech 68235-1-Ig), R-anti-WTAP (1:1000, Abcam ab155655), M-anti-WTAP (1:1000, Proteintech 60188-1-Ig), R-anti-METTL3 (1:1000, Abcam ab195352), M-anti-METTL3 (1:1000, Proteintech, 67733-1-IG), M-anti-NS5A (1:1000, gift of Dr. Charles Rice), R-anti-FLAG (1:500, Sigma Aldrich, F7425), M-anti-FLAG horseradish peroxidase-conjugated (HRP) (1:4000, Sigma Aldrich A8592), M-anti-HA HRP (1:10000, Genetex, GTX18181), G-anti-rabbit Alexa Fluor 568 (1:5000, Thermofisher A-11036).

### Viruses

Infectious stocks of cell culture-adapted strain genotype 2A JFH-1 HCV (JFH-1 M9) were generated, and infectivity was measured by focus-forming assay in Huh7.5 cells as described (Aligeti et al. 2015). For viral infections, cells were incubated in a low volume of serum-free DMEM containing virus at the indicated MOI for 3 h at 37°C. After this time, the cells were replenished with fresh cDMEM. For RIP experiments requiring higher viral RNA input, cells were infected at MOI 1; for phenotypic and meRIP experiments, MOI 0.3 was used.

### Transfection and siRNA Treatment

DNA transfections were performed using FuGENE6 (Promega). siRNA transfection (60 pmol of siRNA) was performed using Lipofectamine RNAiMAX (Invitrogen). Solution was added dropwise to Huh7 cells covered with cDMEM with a media change to cDMEM after 6 hours. The following siRNAs were used in this study: siVIRMA (Horizon: D-019278-02-0005), siMETTL3 (Qiagen: SI04317096), siMETTL14 (Qiagen: SI00459942), siWTAP (Qiagen: SI00069853), and a nontargeting control (siCTRL) (Qiagen: 1027281).

### Cell Lysis

Cells were lysed in either a modified radioimmunoprecipitation assay (RIPA) buffer (TX-100-RIPA) buffer (50 mM Tris [pH 7.5], 150 mM NaCl, 5 mM EDTA, 0.1% sodium dodecyl sulfate (SDS), 0.5% sodium deoxycholate, and 1% Triton X-100), Nonidet P-40 (NP-40) lysis buffer (20 mM Tris [pH 7.4], 100 mM NaCl, and 0.5% NP-40), or Native RIP Buffer (NRB; 10 mM Tris-HCl [pH 7.4], 100 mM KCl, 5 mM MgCl₂, and 0.5% NP-40), each supplemented with protease inhibitor cocktail (Sigma Aldrich, 1:100) and Halt phosphatase inhibitor cocktail (Thermofisher, 1:100). Cells were incubated on ice for 30 minutes and supernatants were collected after centrifugal clarification.

### Immunoblotting

Protein concentrations were determined by Bradford assay (Bio-Rad). Equal amounts of protein were resolved by SDS–polyacrylamide gel electrophoresis using either 4–20% gradient gels (Bio-Rad) with 1X SDS running buffer or 3–8% NuPAGE Tris-acetate gels with 1× NuPAGE Tris-acetate SDS running buffer (Thermofisher). Proteins were transferred to nitrocellulose or polyvinylidene difluoride membranes using Trans-Blot Turbo transfer buffer and the Trans-Blot Turbo transfer system (Bio-Rad). Membranes were stained with REVERT total protein stain (Licor Biosciences) and blocked with 3% bovine serum albumin in 1X Phosphate Buffer Saline with 0.1% Tween (PBS-T). Membranes were probed with primary antibodies directed against proteins of interest, washed with PBS-T, incubated with species specific HRP-conjugated antibodies, washed again with PBS-T, and treated with Clarity Western ECL substrate (Bio-Rad). Imaging was then performed using a LICOR Odyssey FC.

### Immunofluorescence Microscopy

Huh7 cells were fixed and permeabilized in 4% paraformaldehyde in PBS and blocked with 10% FBS in PBS. Slides were stained with indicated primary antibody and washed 3 X in PBS, incubated with conjugated Alexa Fluor secondary antibody, and mounted with ProLong Diamond Antifade Mountant with DAPI (Thermofisher). Imaging was performed on a Leica DM4B widefield fluorescence microscope using a 63x oil objective. All images were processed with National Institutes of Health Fiji/ImageJ.

### Protein Immunoprecipitation

#### For endogenous WTAP IP

Huh7 cells were lysed in NP-40 lysis buffer as described. 500 μg of protein was incubated with 5 μl of R-anti-WTAP antibody coupled to Protein G magnetic beads (Thermofisher) overnight at 4°C in a total volume of 500 μl. Species- and isotype-matched IgG (Cell Signaling) served as the negative control. Beads were then washed with PBS 3 times and eluted in NuPAGE lithium dodecyl sulfate (LDS) sample buffer (Thermofisher) with 1:20 β-mercapto-ethanol (Bio-Rad) and incubated at 70°C for 10 minutes.

#### For IP of transfected proteins

Huh7 cells were lysed in RIPA buffer and prepared as described. 100 μg of protein was incubated with 10 μl of Pierce anti-HA beads (Thermofisher) for 1 h in a total volume of 500 μl. Beads were then washed with PBS 3 times and eluted in 40 μl of 2X Laemmli Sample buffer with 1:20 β-mercaptoethanol (Bio-Rad) and incubated at 95°C for 5 minutes.

### RNA Immunoprecipitation

Huh7 cells or Huh7-K2040 cells were lysed in NRB supplemented with RNasin Plus Ribonuclease Inhibitor (Promega, 1:1000). Cleared lysates were incubated with the indicated antibody-conjugated beads (see below) overnight at 4°C with rotation. Beads were washed five times with NRB. Bound complexes were eluted by proteinase K digestion for 50°C for 30 minutes (Proteinase K (New England Biolabs, 0.3 μg/μl) in 10 mM Tris-Cl [pH 7.4], 100 mM NaCl, 12.5 mM EDTA, 0.1% SDS, supplemented with GlycoBlue Coprecipitant (Thermofisher, 1:1000), and RNasin Plus Ribonuclease Inhibitor (Promega, 1:1000)). RNA was recovered by phenol-chloroform and ethanol precipitation. RNA was reverse-transcribed and quantified by RT-qPCR as described below; enrichment is reported as percent of input. <u>FLAG-VIRMA RIP</u>: Each IP used 300 µg of lysate and 5 µL anti-FLAG magnetic beads (Thermofisher). <u>Endogenous VIRMA RIP</u>: Each IP used 500 µg of lysate and was incubated with 2 µg of VIRMA antibody precoupled to Protein G magnetic beads (Thermofisher); species- and isotype-matched IgG (Cell Signaling) served as the negative control. <u>Endogenous METTL3 RIP</u>: These IPs were performed using the Magna RIP kit protocol (Millipore) according to the manufacturer’s instructions. Each immunoprecipitation used 300 µg of lysate and 3 µg of METTL3 antibody that was precoupled to Protein G magnetic beads; species- and isotype-matched IgG served as the negative control.

### RT-qPCR

Total cellular RNA was extracted from cells using TRIzol (Thermofisher) according to the manufacturer’s protocol. RNA was then reverse transcribed using the iSCRIPT cDNA synthesis kit (Bio-Rad) as per the manufacturer’s instructions. The resulting cDNA was diluted 1:5 in nuclease-free distilled H₂O. RT-qPCR was performed in triplicate using the power SYBR green PCR master mix (Thermofisher) and the Applied Biosystems QuantStudio 6 Flex RT-PCR system. qPCR technical replicates were assessed for consistency prior to analysis. Ct values with SD >0.5 across three technical replicates were flagged, and outlier Ct values >0.5 from the median were excluded, retaining a minimum of two technical replicates per biological sample. This analysis was applied uniformly across all targets, conditions, and biological replicates prior to fold-change calculation. Primer sequences for RT-qPCR are listed in Data Table S1.

### MeRIP-qPCR

meRIP was performed as previously described on total or fragmented RNA (NEB, EpiMark® N6-Methyladenosine Enrichment Kit) (McFadden et al. 2021). All experiments were carried out using 15-20 µg of total RNA per IP condition. Following meRIP, cDNA from the input and immunoprecipitated RNA fractions was generated and analyzed by RT-qPCR. The relative m^6^A level for each transcript was calculated as the percentage of input under each condition relative to siCTRL or vector. Controls for meRIP specificity include an unmodified RNA spike-in and m^6^A-modified spike-in, which are added to each sample prior to immunoprecipitation. Recovery of the spike-in RNAs was quantified by RT-qPCR (**Figure S1**).

### Statistical Analysis

Statistical analysis was performed using GraphPad Prism 9. Data-appropriate statistical tests were performed, including Welch’s t-test and 1-way ANOVA with post-hoc testing. Values are presented as mean ± standard error of the mean for biological replicates (n=3, or as indicated). *-P < 0.05, ** - P < 0.01, *** - P < 0.001.

### Declaration of Generative AI in the Writing Process

During the preparation of this work, the authors used Perplexity (including its Deep Research and Orchestrator features) and ChatGPT (OpenAI) to improve readability and language. After using these tools, the authors reviewed and edited the content as needed and take full responsibility for the content of the publication.

## REFERENCES

1. Aligeti M, Roder A, Horner SM. 2015. Cooperation between the Hepatitis C Virus p7 and NS5B Proteins Enhances Virion Infectivity. J Virol 89: 11523–11533.

2. Arguello AE, DeLiberto AN, Kleiner RE. 2017. RNA Chemical Proteomics Reveals the N(6)-Methyladenosine (m(6)A)-Regulated Protein-RNA Interactome. J Am Chem Soc 139: 17249–17252.

3. Baquero-Perez B, Geers D, Diez J. 2021. From A to m(6)A: The Emerging Viral Epitranscriptome. Viruses 13: 1049.

4. Bawankar P, Lence T, Paolantoni C, Haussmann IU, Kazlauskiene M, Jacob D, Heidelberger JB, Richter FM, Nallasivan MP, Morin V et al. 2021. Hakai is required for stabilization of core components of the m(6)A mRNA methylation machinery. Nat Commun 12: 3778.

5. Bermudez Y, Hatfield D, Muller M. 2024. A Balancing Act: The Viral-Host Battle over RNA Binding Proteins. Viruses 16.

6. Castello A, Kamel W. 2025. Nuclear RNA-binding proteins meet cytoplasmic viruses. RNA 31: 444–451.

7. Edupuganti RR, Geiger S, Lindeboom RGH, Shi H, Hsu PJ, Lu Z, Wang SY, Baltissen MPA, Jansen P, Rossa M et al. 2017. N(6)-methyladenosine (m(6)A) recruits and repels proteins to regulate mRNA homeostasis. Nat Struct Mol Biol 24: 870–878.

8. Filbin ME, Kieft JS. 2009. Toward a structural understanding of IRES RNA function. Curr Opin Struct Biol 19: 267–276.

9. Filbin ME, Vollmar BS, Shi D, Gonen T, Kieft JS. 2013. HCV IRES manipulates the ribosome to promote the switch from translation initiation to elongation. Nat Struct Mol Biol 20: 150–158.

10. Garcias Morales D, Reyes JL. 2021. A birds’-eye view of the activity and specificity of the mRNA m(6) A methyltransferase complex. Wiley Interdiscip Rev RNA 12: e1618.

11. Gilbert WV, Nachtergaele S. 2023. mRNA Regulation by RNA Modifications. Annu Rev Biochem 92: 175–198.

12. Gokhale NS, McIntyre ABR, Mattocks MD, Holley CL, Lazear HM, Mason CE, Horner SM. 2020. Altered m(6)A Modification of Specific Cellular Transcripts Affects Flaviviridae Infection. Mol Cell 77: 542–555 e548.

13. Gokhale NS, McIntyre ABR, McFadden MJ, Roder AE, Kennedy EM, Gandara JA, Hopcraft SE, Quicke KM, Vazquez C, Willer J et al. 2016. N6-Methyladenosine in Flaviviridae Viral RNA Genomes Regulates Infection. Cell Host Microbe 20: 654–665.

14. Horner SM, Reaves JV. 2024. Recent insights into N(6)-methyladenosine during viral infection. Curr Opin Genet Dev 87: 102213.

15. Iselin L, Palmalux N, Kamel W, Simmonds P, Mohammed S, Castello A. 2022. Uncovering viral RNA-host cell interactions on a proteome-wide scale. Trends Biochem Sci 47: 23–38.

16. Kim GW, Imam H, Khan M, Siddiqui A. 2020. N(6)-Methyladenosine modification of hepatitis B and C viral RNAs attenuates host innate immunity via RIG-I signaling. J Biol Chem 295: 13123–13133.

17. Kim GW, Siddiqui A. 2021. N6-methyladenosine modification of HCV RNA genome regulates cap-independent IRES-mediated translation via YTHDC2 recognition. Proc Natl Acad Sci U S A 118.

18. Lee Q, Song R, Phan DAV, Pinello N, Tieng J, Su A, Halstead JM, Wong ACH, van Geldermalsen M, Lee BS et al. 2023. Overexpression of VIRMA confers vulnerability to breast cancers via the m(6)A-dependent regulation of unfolded protein response. Cell Mol Life Sci 80: 157.

19. Li N, Zhu Z, Deng Y, Tang R, Hui H, Kang Y, Rana TM. 2023. KIAA1429/VIRMA promotes breast cancer progression by m(6) A-dependent cytosolic HAS2 stabilization. EMBO Rep 24: e55506.

20. Lindenbach BD, Rice CM. 2005. Unravelling hepatitis C virus replication from genome to function. Nature 436: 933–938.

21. Liu J, Yue Y, Han D, Wang X, Fu Y, Zhang L, Jia G, Yu M, Lu Z, Deng X et al. 2014. A METTL3-METTL14 complex mediates mammalian nuclear RNA N6-adenosine methylation. Nat Chem Biol 10: 93–95.

22. McFadden MJ, McIntyre ABR, Mourelatos H, Abell NS, Gokhale NS, Ipas H, Xhemalce B, Mason CE, Horner SM. 2021. Post-transcriptional regulation of antiviral gene expression by N6-methyladenosine. Cell Rep 34: 108798.

23. Moradpour D, Penin F, Rice CM. 2007. Replication of hepatitis C virus. Nat Rev Microbiol 5: 453–463.

24. N’Da Konan S, Segeral E, Bejjani F, Bendoumou M, Ait Said M, Gallois-Montbrun S, Emiliani S. 2022. YTHDC1 regulates distinct post-integration steps of HIV-1 replication and is important for viral infectivity. Retrovirology 19: 4.

25. Paul D, Bartenschlager R. 2015. Flaviviridae Replication Organelles: Oh, What a Tangled Web We Weave. Annu Rev Virol 2: 289–310.

26. Ping XL, Sun BF, Wang L, Xiao W, Yang X, Wang WJ, Adhikari S, Shi Y, Lv Y, Chen YS et al. 2014. Mammalian WTAP is a regulatory subunit of the RNA N6-methyladenosine methyltransferase. Cell Res 24: 177–189.

27. Ramage HR, Kumar GR, Verschueren E, Johnson JR, Von Dollen J, Johnson T, Newton B, Shah P, Horner J, Krogan NJ et al. 2015. A combined proteomics/genomics approach links hepatitis C virus infection with nonsense-mediated mRNA decay. Mol Cell 57: 329–340.

28. Romero-Brey I, Merz A, Chiramel A, Lee JY, Chlanda P, Haselman U, Santarella-Mellwig R, Habermann A, Hoppe S, Kallis S et al. 2012. Three-dimensional architecture and biogenesis of membrane structures associated with hepatitis C virus replication. PLoS Pathog 8: e1003056.

29. Sacco MT, Bland KM, Horner SM. 2022. WTAP Targets the METTL3 m(6)A-Methyltransferase Complex to Cytoplasmic Hepatitis C Virus RNA to Regulate Infection. J Virol 96: e0099722.

30. Sarnow P, Sagan SM. 2016. Unraveling the Mysterious Interactions Between Hepatitis C Virus RNA and Liver-Specific MicroRNA-122. Annu Rev Virol 3: 309–332.

31. Schwartz S, Mumbach MR, Jovanovic M, Wang T, Maciag K, Bushkin GG, Mertins P, Ter-Ovanesyan D, Habib N, Cacchiarelli D et al. 2014. Perturbation of m6A writers reveals two distinct classes of mRNA methylation at internal and 5’ sites. Cell Rep 8: 284–296.

32. Slobodin B, Han R, Calderone V, Vrielink J, Loayza-Puch F, Elkon R, Agami R. 2017. Transcription Impacts the Efficiency of mRNA Translation via Co-transcriptional N6- adenosine Methylation. Cell 169: 326–337 e312.

33. Srinivas KP, Depledge DP, Abebe JS, Rice SA, Mohr I, Wilson AC. 2021. Widespread remodeling of the m(6)A RNA-modification landscape by a viral regulator of RNA processing and export. Proc Natl Acad Sci U S A 118.

34. Su S, Li S, Deng T, Gao M, Yin Y, Wu B, Peng C, Liu J, Ma J, Zhang K. 2022. Cryo-EM structures of human m(6)A writer complexes. Cell Res 32: 982–994.

35. Sumpter R, Jr., Loo YM, Foy E, Li K, Yoneyama M, Fujita T, Lemon SM, Gale M, Jr. 2005. Regulating intracellular antiviral defense and permissiveness to hepatitis C virus RNA replication through a cellular RNA helicase, RIG-I. J Virol 79: 2689–2699.

36. Sumpter R, Jr., Wang C, Foy E, Loo YM, Gale M, Jr. 2004. Viral evolution and interferon resistance of hepatitis C virus RNA replication in a cell culture model. J Virol 78: 11591–11604.

37. Wan H, Adams RL, Lindenbach BD, Pyle AM. 2022. The In Vivo and In Vitro Architecture of the Hepatitis C Virus RNA Genome Uncovers Functional RNA Secondary and Tertiary Structures. J Virol 96: e0194621.

38. Wang X, Zhao BS, Roundtree IA, Lu Z, Han D, Ma H, Weng X, Chen K, Shi H, He C. 2015. N(6)- methyladenosine Modulates Messenger RNA Translation Efficiency. Cell 161: 1388–1399.

39. Wang Y, Li Y, Toth JI, Petroski MD, Zhang Z, Zhao JC. 2014. N6-methyladenosine modification destabilizes developmental regulators in embryonic stem cells. Nat Cell Biol 16: 191–198.

40. World Health Organization. 2026. Global hepatitis report 2026. World Health Organization, Geneva.

41. Yan X, Pei K, Guan Z, Liu F, Yan J, Jin X, Wang Q, Hou M, Tang C, Yin P. 2022. AI-empowered integrative structural characterization of m(6)A methyltransferase complex. Cell Res 32: 1124–1127.

42. Yue Y, Liu J, Cui X, Cao J, Luo G, Zhang Z, Cheng T, Gao M, Shu X, Ma H et al. 2018. VIRMA mediates preferential m(6)A mRNA methylation in 3’UTR and near stop codon and associates with alternative polyadenylation. Cell Discov 4: 10.

43. Zhang K, Zhang Y, Maharjan Y, Sugiokto FG, Wan J, Li R. 2021. Caspases Switch off the m(6)A RNA Modification Pathway to Foster the Replication of a Ubiquitous Human Tumor Virus. mBio 12: e0170621.

